# Topography of speech-related acoustic and phonological feature encoding throughout the human core and parabelt auditory cortex

**DOI:** 10.1101/2020.06.08.121624

**Authors:** Liberty S. Hamilton, Yulia Oganian, Edward F. Chang

## Abstract

Speech perception involves the extraction of acoustic and phonological features from the speech signal. How those features map out across the human auditory cortex is unknown. Complementary to noninvasive imaging, the high spatial and temporal resolution of intracranial recordings has greatly contributed to recent advances in our understanding. However, these approaches are typically limited by piecemeal sampling of the expansive human temporal lobe auditory cortex. Here, we present a functional characterization of local cortical encoding throughout all major regions of the primary and non-primary human auditory cortex. We overcame previous limitations by using rare direct recordings from the surface of the temporal plane after surgical microdissection of the deep Sylvian fissure between the frontal and temporal lobes. We recorded neural responses using simultaneous high-density direct recordings over the left temporal plane and the lateral superior temporal gyrus, while participants listened to natural speech sentences and pure tone stimuli. We found an anatomical separation of simple spectral feature tuning, including tuning for pure tones and absolute pitch, on the superior surface of the temporal plane, and complex tuning for phonological features, relative pitch and speech amplitude modulations on lateral STG. Broadband onset responses are unique to posterior STG and not found elsewhere in auditory cortices. This onset region is functionally distinct from the rest of STG, with latencies similar to primary auditory areas. These findings reveal a new, detailed functional organization of response selectivity to acoustic and phonological features in speech throughout the human auditory cortex.

## Introduction

Spoken language contains a myriad of acoustic cues that give rise to the rich experience of speech. Listening to speech activates populations of neurons in multiple functional regions of the auditory cortex, including primary and nonprimary cortical fields. The computations carried out in these areas are thought to be responsible for extracting meaningful linguistic content, such as consonants and vowels, or prosodic rhythm and intonation, from the spectrotemporal acoustics of speech. However, the precise feature representations for speech and the relationship between anatomical and functional boundaries in human auditory cortex are disputed.

The majority of previous studies used functional neuroimaging (e.g. fMRI and MEG) to show a progressive selectivity for more complex sound types from primary (medial) towards higher-order (lateral) areas (Brodbeck et al., 2018; Chevillet et al., 2011; De Heer et al., 2017; Rauschecker and Scott, 2009; Wessinger et al., 2001), finding stronger responses to complex stimuli, such as speech and music than simple artificial stimuli, such as pure tones, in higher-order areas (Leaver and Rauschecker, 2010; Norman-Haignere et al., 2015; Schönwiesner and Zatorre, 2009). These results are typically interpreted in the framework of an anatomical-hierarchical model, in which simple acoustic features are transformed into more complex categorical representations across the auditory cortex. Yet, the specific nature of speech sound representations in these areas remain highly controversial. The limited spatial or temporal resolution of non-invasive imaging precludes mapping of locally heterogeneous neural response types, leaving fundamental questions about speech sound representations unanswered. In particular, where and which specific acoustic or linguistic speech features are encoded across the auditory cortex is not known.

Understanding the nature of local sound representations has been greatly aided by the high spatiotemporal resolution of intracranial recordings (Chang, 2015; Lachaux et al., 2012; Flinker et al. 2011; Nourski et al. 2012; Howard et al. 2000; Berezutskaya et al. 2017). The human auditory cortex lies across an expansive surface of the temporal lobe, including the superior temporal plane and the adjacent superior temporal gyrus (STG, Figure 1) on its lateral side. STG is exposed on the lateral temporal lobe and thus accessible by surface electrocorticography (ECoG) recording methods. In contrast, most of the temporal plane, including the tonotopic “auditory core” on posteromedial portion of Heschl’s gyrus, is buried deep within the lateral fissure, which separates the temporal lobe from the frontal and parietal lobes (Brewer and Barton, 2016). These anatomical constraints make the temporal plane difficult to access. Therefore, functional parcellations beyond the core and their relation to anatomical landmarks are not well understood.

**Figure 1.**
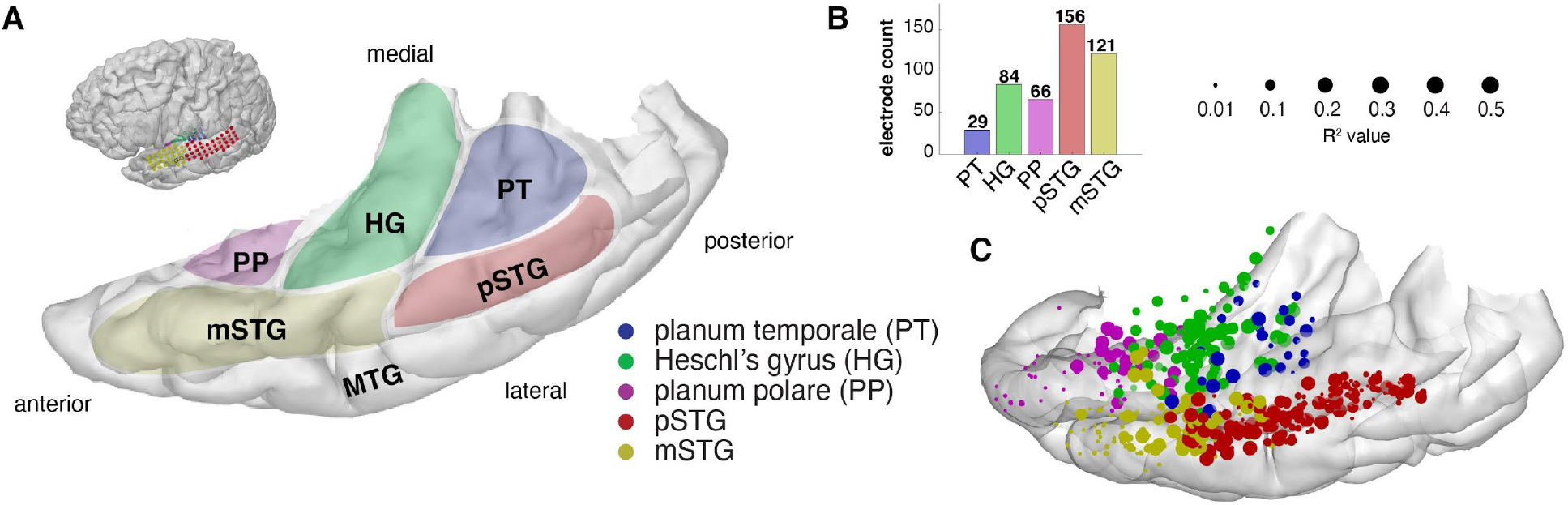
Anatomical parcellations of temporal lobe regions of the human auditory cortex and electrode coverage. (A) Anatomical regions of interest on the left hemisphere temporal lobe of an example participant. Planum temporale (PT) is located posterior to the Heschl’s gyrus (HG), which includes either one or two transverse temporal gyri and is thought to contain the auditory “core” posteromedially. Planum polare (PP) is the anatomical region on top of the temporal plane that is anterior to HG. On the lateral surface, we defined posterior superior temporal gyrus (pSTG) as any region of the superior temporal gyrus that was posterior to the location at which the posterior border of the HG (transverse temporal sulcus) exits to the lateral surface of the STG. Middle STG (mSTG) was defined as the anatomical region anterior to this border. Middle temporal gyrus is located inferiorly to the STG. (B) Electrode counts across anatomical areas for all 9 participants. (C) All participants’ electrodes projected onto an MNI atlas brain (cvs_avg35_inMNI152). Electrode size reflects the maximum amount of variance (R^2^) explained by the encoding models tested in our analyses, across different types of represented information. Electrode sites are colored according to their anatomical location on PT, HG, PP, pSTG or mSTG. Local overlap between colors reflects variability in warping of individual brain coordinates to the MNI space, as electrode location was determined based on individual brain anatomy.

Several studies use an encoding model framework, in which different parameterizations of the stimulus (e.g. the spectrogram, pitch contour, phoneme timings, or phonological feature timings) are compared to determine which representation best predicts the observed neural data (Aertsen and Johannesma, 1981; De Heer et al., 2017; Holdgraf et al., 2017; Theunissen et al., 2001). Our recent work using intracranial electrocorticography (ECoG) with high density grids and widespread sampling of the lateral STG have revealed a patchwork of specialized neural populations selective for different speech features, including phonological features (Chang et al., 2010; Mesgarani et al., 2014), speech-range spectrotemporal modulations (Hullett et al., 2016), acoustic onsets (Hamilton et al., 2018), syllabic nucleus onsets (Oganian and Chang, 2019), and intonational prosody and pitch (Tang et al., 2017). In contrast, recordings from medial temporal areas typically use penetrating depth electrodes, which may capture the long axis of Heschl’s gyrus, but do not have the uniform and widespread cortical surface coverage offered by electrode grids (Berezutskaya et al., 2017; Brugge et al., 2003; Griffiths et al., 2010; Nourski et al., 2014; Steinschneider et al., 2014). In addition, few studies also record from the planum temporale and planum polare, so these areas remain relatively understudied (Besle et al., 2008; Bidet-Caulet et al., 2007; Liégeois-Chauvel et al., 1999). Placing grid electrodes in this area requires meticulous surgical dissection of the Sylvian fissure and is only possible in cases of opercular/insular surgery (Bouthillier et al., 2012; Malak et al., 2009).

Overall, while recent advances in human intracranial recordings have revealed specialization for different speech sound features in lateral STG, whether similar feature representations exist on the superior temporal surface, including core auditory cortex, and whether STG responses to non-speech reflect the same computation principles remain open questions. Such an analysis requires sampling of neural responses to natural speech and experimentally controlled simple sound stimuli simultaneously across all cortical auditory areas. Critically, a comprehensive map of feature encoding across primary and higher-order auditory areas is prerequisite to a meaningful evaluation of models of information flow and transformations of cortical representations.

Here we aim to address this by mapping out which speech sound features are represented in different regions of the auditory cortex. We used pure tone and natural speech stimuli to address the dependence of neural responses on stimulus complexity and to bridge between studies that focused on solely one type of stimulus or a subset of auditory fields. Using high-density electrode grids, we simultaneously recorded neural activity from multiple subfields of the human temporal lobe auditory cortex. The anatomical divisions of the human auditory cortex include the planum temporale (PT), Heschl’s gyrus (HG; or transverse temporal gyrus), and planum polare (PP) on the superior temporal plane (Moerel et al., 2014), and posterior (pSTG) and middle superior temporal gyrus (mSTG) on its lateral surface (Figure 1A). Here, we define pSTG as the portion of the STG posterior to the lateral exit point of the transverse temporal sulcus (Friederici, 2015; Upadhyay et al., 2008). The high spatial and temporal resolution of our recordings allowed us to assess the intersection of functional and anatomical organization within the human auditory cortex, incorporating feature representations, temporal response dynamics, and functional parcellations.

## Results

We acquired direct intracranial recordings from 636 electrode sites across the left temporal plane and STG in 9 participants, including both grid and depth electrodes (Figure 1B, Figure S1), as participants listened to speech and pure-tone stimuli. The distribution of speech-selective electrodes across auditory regions is shown in Figure 1B, where electrodes are colored according to the percent variance explained by the best linear receptive field model in the speech data (see Methods). Electrodes on each participant’s brain are shown in Figure S1.

### Preferential tuning to pure tones on the superior temporal plane and to speech in STG

To investigate tuning for simple spectral versus complex speech-specific features across the anatomical regions comprising the auditory cortex, we played pure tone and natural sentence stimuli to participants while simultaneously recording local field potentials from the temporal plane and lateral superior temporal gyrus. All patients (N=9) heard the speech stimuli, which included 499 sentences from the TIMIT acoustic-phonetic database (Garofolo et al., 1993) spoken by male and female talkers. Due to timing constraints in the intraoperative setting, a subset of patients (N=5) were presented with pure tone stimuli at three amplitude levels and a frequency range that matched speech stimuli (0.75 Hz to 8 kHz, see Methods). Short pure tone stimuli similar to those used here are typically used to identify tonotopic gradients associated with auditory “core” regions, as they robustly activate frequency-specific regions across the auditory pathway (Brewer and Barton, 2016; Moerel et al., 2012; Wessinger et al., 1997, 2001). Using both types of stimuli allowed us to compare the representations of simple and complex auditory stimuli throughout the auditory cortical hierarchy.

Our first question was whether tones and speech would induce neural responses of comparable magnitude across areas (Fig 2B). Tone response magnitude was measured as the peak high gamma response across all tone stimuli, and speech response magnitude was measured as the peak high gamma response across all sentence stimuli. Overall, we found neural responses to tone and speech stimuli throughout all areas. We observed significant responses to tones as compared to silence in 100% of PT sites, 93% of HG, 51% of PP, 70% of pSTG sites, and 61% of mSTG sites (Bonferroni-corrected p<0.05, Wilcoxon rank sum test). For the same electrodes used in this tone analysis, we observed significant responses to speech in 100% of PT electrodes, 100% of HG electrodes, 77% of PP electrodes, 86% of pSTG electrodes, and 81% of mSTG electrodes. Only 6.4% of electrodes showed significant responses to tones and not speech, 60% of electrodes showed a significant response to both speech and tones, and of note, 24% showed a significant response to speech but not to tones. Of the population of electrodes selective to speech and not tones, the vast majority were located outside of the putative auditory core (0% were in PT, 2% HG, 33% in PP, 35% in pSTG, and 30% in mSTG).

**Figure 2.**
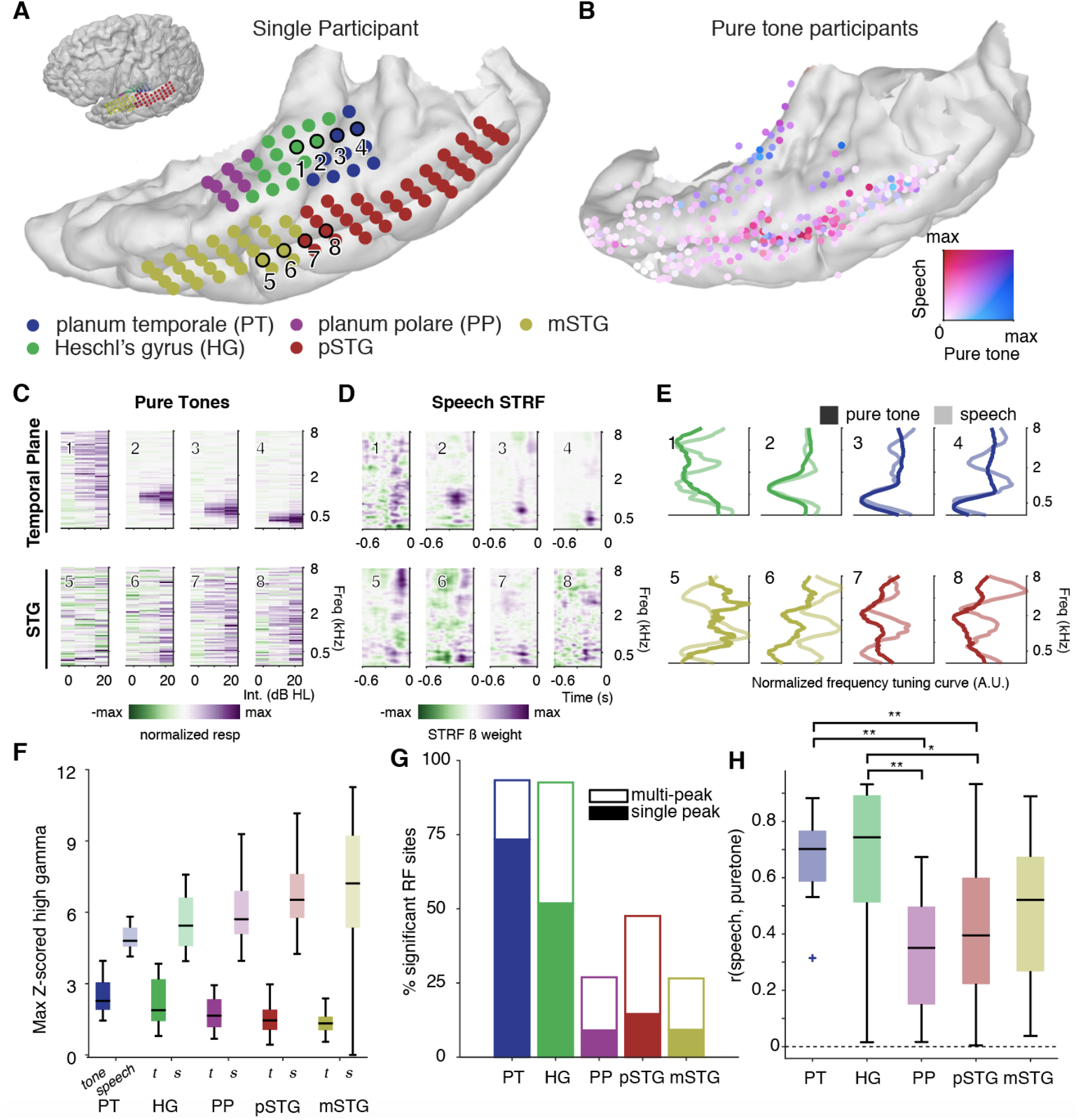
Regional selectivity for speech and pure tones; divergence of tuning curves to simple and complex acoustic inputs in the human auditory cortex. (A) Example temporal plane and lateral temporal cortical grids on one participant’s temporal lobe reconstruction. Electrodes are colored as in A. Small inset shows the position of the temporal lobe in the whole brain reconstruction. (B) Comparison between normalized response magnitudes to speech and pure tone stimuli across all participants, superimposed on the MNI brain. Electrodes are colored according to the normalized magnitude of the pure tone response (blue) and speech response during sentence listening (red). Mixed speech/pure tone responses are shown in purple. (C) Example pure tone receptive fields for electrodes shown in A. (D) Example speech spectrotemporal receptive fields (STRFs) for electrodes in A. (E) Comparison of normalized pure tone and speech tuning curves from sites in C and D. Response curves were normalized for each stimulus and electrode separately to focus the figure on relative response magnitudes across frequency bands. (F) Maximum pure tone response and speech responses by anatomical area. Pure tone response magnitudes dropped off in more lateral sites and significantly differed across anatomical regions (Kruskal Wallis ANOVA, Chi-sq=34.39, df=4, p=6.2×10-7). Speech response magnitudes increased toward more lateral sites (Kruskal Wallis ANOVA, Chi-Sq=46.8, df=4, p=1.7×10-9). (G) Percentage of sites with significant receptive fields by anatomical area, as measured by significance of within RF responses as compared to outside RF. Percentages are split into single-versus multi-peaked RF. (H) Correlation between frequency tuning for pure tones (as in D) to frequency tuning for speech (from STRF, as in E). Responses in PT and HG are most similar for pure tones and speech. For all boxplots, boxes indicate the 25th and 75th percentiles and the median. Whiskers show extreme non-outlier values.

Although the anatomical areas within auditory cortex mostly showed responses to both classes of sound, they differed in the magnitude of their responses, reflected in opposing gradients for speech and tone response magnitudes from medial to lateral areas (see Table 2 for all pairwise comparisons between anatomical areas). That is, responses to speech increased from medial to lateral regions, whereas responses to tones decreased. Responses to tones were stronger in PT and HG as compared to STG (Figure 2B, 2F). In the example shown in Figure 1C, neural responses from adjacent electrodes demonstrated tonotopic organization of the ordered selectivity to sound frequency. Pure tone receptive fields for all recorded sites are shown in Figures S2 and S3. In contrast, sites on STG responded more strongly to speech than sites on the superior temporal plane (Fig 2B-D, 2F). This difference in selectivity is consistent with previous reports of selective preference for complex/speech stimuli in STG and strong tuning to pure frequency tones on the superior temporal plane (Binder et al., 2000; Démonet et al., 1992; Leaver and Rauschecker, 2010, 2016; Nourski et al., 2012; Steinschneider et al., 2013, 2014).

In addition to the overall differences in response magnitude to pure tones and speech, we quantified classical tone frequency receptive field (RF) shape and quality (Figure 2G). To determine whether each site had a strongly tuned classical RF, we divided the entire tone frequency range into ‘inside’ and ‘outside’ of the RF using a binary mask (see Methods). Frequency tuning was significant if responses to tones inside the classical RF were significantly higher than responses to tones outside the RF (Wilcoxon rank sum test). This analysis distinguishes between nonselective overall changes in baseline activity during tone presentation (e.g. example electrodes 5-8 in Figure 2E) versus clear V-shaped classical receptive fields where the tone response is clearly isolated to specific frequency-intensity pairs (Figure 2E electrodes 1-4). For these clear V-shaped RFs, we also distinguished between single-peaked receptive fields (for example electrodes 2-4 in Figure 2E) versus multi-peaked (≥2 peaks, for example electrode 1 in Figure 2E). We expected to see strong, single-peaked frequency tuning in putative auditory core areas, with broader and more diffuse tuning in lateral and anterior sites. As expected, we found significant tone selectivity with V-shaped tuning curves on the vast majority of PT (93%) and HG (93%) sites, with most of these sites showing well-defined single-peaked RFs (79% for PT and 56% for HG respectively). Anterior sites in PP, however, were much less likely to have significant frequency tuning of their receptive fields (27%), and were more likely to be spectrally broad and multi-peaked (33% single-peaked). Lateral sites in pSTG and mSTG also showed a lower proportion of clear V-shaped RFs compared to HG and PT (pSTG = 48%, with 31% single-peaked and mSTG = 27%, with 65% single-peaked). The proportion of non-tone responsive, single-, and multi-peaked RFs differed significantly across all five anatomical areas (chi^2^=89.9, df=8, N=342 electrodes, p=4.8×10^−16^).

### Tuning for pure tones does not predict responses to speech outside of the core

We next asked whether frequency tuning curves were similar for speech, since previous studies have shown that responses to simple, synthesized stimuli may not predict responses to more complex, natural stimuli (Hamilton and Huth, 2018; Hullett et al., 2016; Theunissen et al., 2000, 2001; Rauschecker and Tian, 2000). By directly comparing responses to pure tones and TIMIT sentences in the same electrodes, we found a significant difference between core and the lateral auditory areas.

Visual inspection showed that in the narrowly tuned sites of the auditory core, frequency tuning for tones and speech was similar (Fig. 2C-D, top panels), but in many sites outside of HG and PT core auditory cortex, speech and pure tone tuning diverged (Fig. 2C-D, bottom panels). To address this, we calculated the Pearson correlation between the frequency tuning curve assessed from the pure tone response (as in Fig. 2C) and the spectrotemporal receptive field (STRF) analysis in response to speech (as in Fig. 2D). Frequency tuning curves were calculated by collapsing across intensities (for pure tones) and time delays (for speech STRFs) and are depicted in Fig. 2E for example electrodes. Electrodes with a similar pure tone and speech response (e.g. electrode 2 in Fig. 2E) would appear more correlated in this analysis. Conversely, electrodes with different pure tone and speech responses would show a low correlation (e.g. electrode 5 in Fig. 2E).

Frequency tuning from speech and tones was highly similar in PT and HG, but diverged for other areas (Fig. 2H). The correlation between pure tone and speech frequency tuning was significantly different across anatomical regions (Kruskal-Wallis ANOVA, Chi-sq=23.01, df=4, p=0.0001). Post-hoc tests using the Tukey-HSD correction for multiple comparisons showed that correlations were significantly higher in PT compared to PP (p=0.005) and PP compared to pSTG (p=0.003). Correlations were also higher in HG compared to PP (p=0.0009) and pSTG (p=0.01). Notably, this performance difference was not due to lower performance of the spectro-temporal model for speech in lateral STG regions (Figure 1B) -- in addition, only electrodes with R^2^-values > 0.01 for speech STRFs or significant tone responses were included in this analysis. Both PT and HG also had the highest percentage of sites with correlations between the speech and pure tone tuning curves that were significantly greater than 0 (PT: 82%, HG: 71%. PP: 36%, pSTG: 31%, mSTG: 33%). These findings demonstrate that speech responses in the core areas can be well predicted from their responses to pure tones, whereas in lateral STG and other non-core areas they cannot and are likely tuned to more complex combinations of spectral features.

### Functional organization of speech feature encoding throughout the auditory cortex

The difference in spectral tuning as estimated from tones and speech suggests that non-primary areas represent complex acoustic patterns. We thus turned to characterize the encoding of acoustic and phonological features across different areas of human auditory cortices. We expected that neural responses in more primary areas on the superior temporal plane may be best captured by a spectrotemporal model, preserving the detailed characteristics of sound acoustics, but without additional assumptions about the linguistic content of the stimulus. In contrast, we expected that on lateral STG, neural responses would represent specific linguistic features while remaining invariant to other acoustic characteristics. For example, previous work showed separate neural populations that tracked the relative pitch contour of a speech stimulus regardless of phoneme content, as well as the opposite -- regions that responded equally to the same phoneme regardless of pitch (Tang et al., 2017). Regions that are purely spectrotemporal might respond to a particular vowel, but would respond differently depending on whether that vowel was spoken by a person with a high-pitched versus a low-pitched voice. Finally, we wanted to determine whether responses we observed previously for sound onsets (Hamilton et al., 2018) and envelope change (Oganian and Chang, 2019) on lateral STG were unique, or also present on the temporal plane.

We thus fit models describing tuning for spectrotemporal features (Figure 2D) as well as mixed feature representations including the temporal structure of speech (speech onsets from silence and peaks in the rate of loudness change, peakRate), phonological features, absolute pitch, and relative pitch (speaker-normalized relative pitch and relative pitch rate of change), as depicted in Figure 3A. Phonological features included more global “sonorant” versus “obstruent” categories, which distinguish roughly between vowels and consonants based on the obstruction of airflow in the vocal tract, and also categories for plosive, fricative, or high/front/low/back vowels, which describe in more detail how a speech sound is made based on the manner or place of articulation. By stepwise extending models to include additional features and comparing resulting model R^2^, we captured the relative contributions of each set of acoustic-phonetic features. This analysis revealed that neural populations captured by single electrodes selectively encode only some but not other features in the speech signal. Figure 3B shows receptive fields of the best encoding models for five example electrodes in different participants: E1 and E2 encode mainly the temporal cues of onset and peakRate, respectively, and E3 also represents sonorant, voiced, and high vowel phonological features. In contrast, absolute pitch features explained a significant amount of variance in E4, while E5 encoded relative pitch features. We observed a wide diversity of responses across the human auditory cortex.

**Figure 3.**
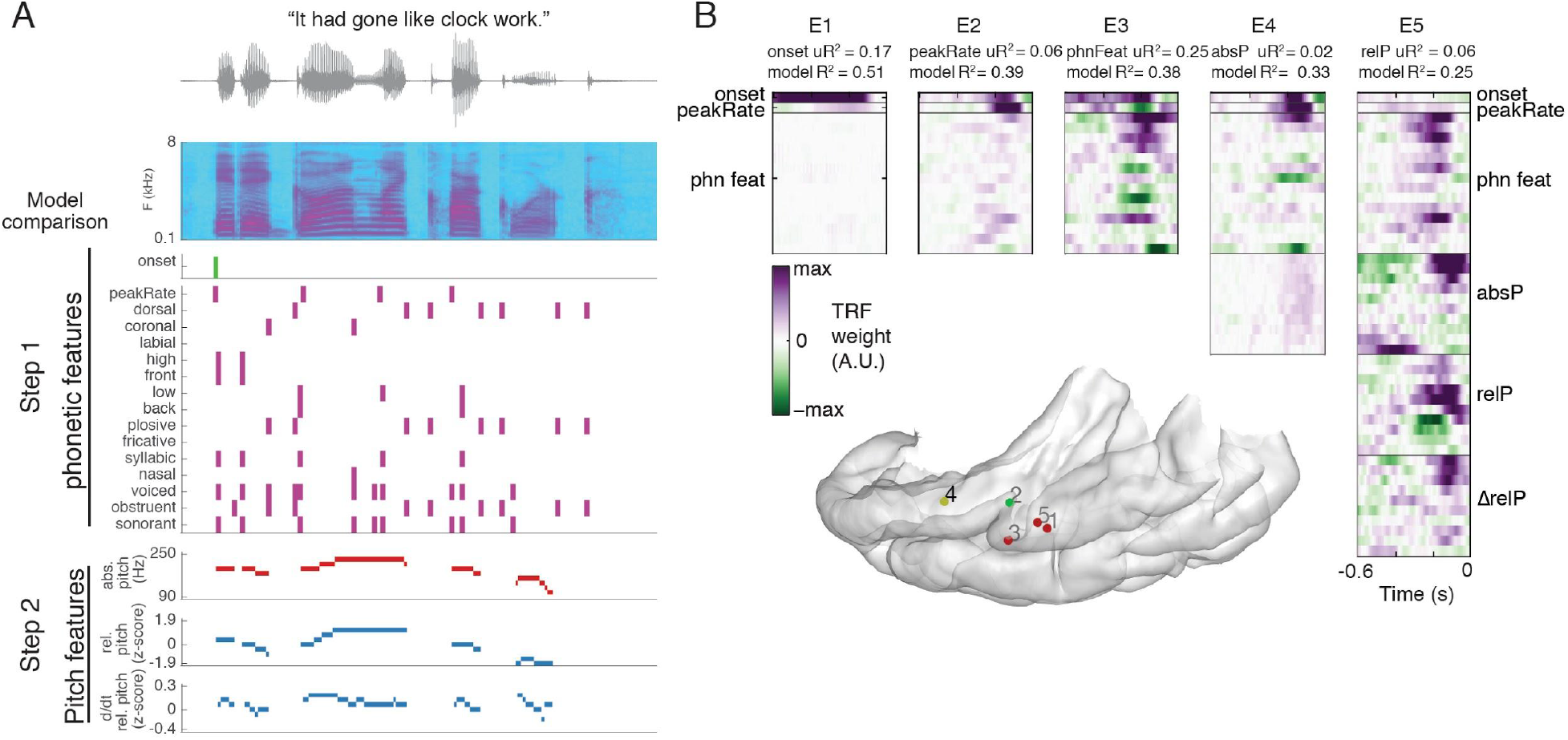
Speech features and example receptive fields. **(A)** Speech features tested in feature model comparisons for an example sentence. **(B)** Best model receptive fields for example electrodes. For each electrode, the best-performing model is plotted. Electrodes were chosen for which distinct sets of features explain a large portion of variance. For example, for electrode 1, the best model includes onset, peakRate and phonetic features, but 17% of variance are attributed to the onset predictor.

This analysis revealed a functional and anatomical localization of onset encoding in pSTG, peakRate, feature and relative pitch encoding in mid-STG, and absolute pitch encoding in planum temporale and Heschl’s gyrus (Figure 4A, 4B). First, we compared the full spectro-temporal model to smaller models that contains binary speech features, but no information about the precise spectro-temporal structure of the speech stimulus. Comparing a reduced model containing only an onset predictor to a full spectro-temporal model, we found that the additional spectro-temporal resolution did not improve model fits for a subset of electrodes, located in posterior STG (Figure 4C, F). Second, a similar comparison showed that for a group of electrodes in middle STG, a phonological feature model (including onsets, peakRate and phonological features in 15 predictors with no specific spectro-temporal detail) outperformed the full spectro-temporal model (Figure 4B, E). We combined phonological features and peakRate together in this step, as more detailed model comparisons showed these features were represented in mostly overlapping sets of electrodes, in line with our previous work (Oganian and Chang, 2019). In particular, we found that both peakRate and phonetic features were encoded predominantly in middle STG (Fig. 4 C, E). The better performance of a reduced set of features over the full spectro-temporal description of the acoustic stimulus in the lateral STG are in line with our previous work (Hamilton et al., 2018; Mesgarani et al., 2014). The confinement of such electrodes to the lateral STG (Fig. 4A) supports the unique role of lateral STG in speech processing.

**Figure 4.**
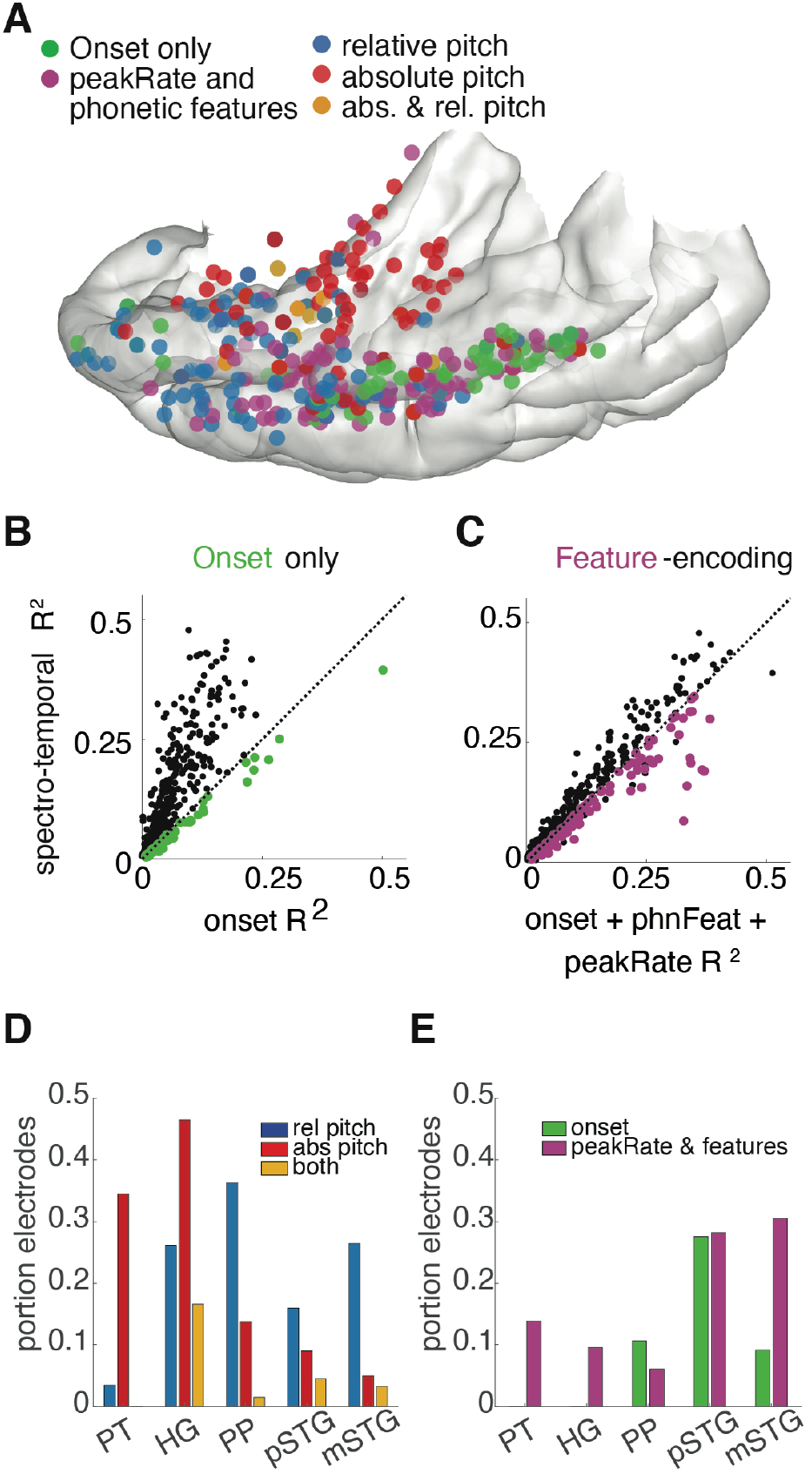
Anatomical separation of selectivity for speech features. (A) Anatomical location of electrodes primarily coding for speech onsets, phonetic features and peakRate, relative pitch and absolute pitch. (B) Onset and (C) feature-encoding electrodes are defined as those for which the respective model outperforms a spectro-temporal model. (D) Anatomical distribution of pitch encoding. Absolute pitch is primarily represented on temporal plane, relative pitch is primarily represented in planum polare and anterior STG (chi-sq = 62.5, p < 10-9). [both = both rel and abs pitch contribute unique variance, permutation p<.05] Proportion electrodes relative to number of speech-responsive electrodes in each anatomical area. (E) Anatomical distribution of electrode encoding onsets and phonological features+peakRate. Onset-encoding electrodes are mostly located in pSTG, whereas phonological features and peakRate are represented in posterior and anterior lateral STG. Proportion electrodes relative to number of speech-responsive electrodes in each anatomical area.

In the next step, to identify encoding of pitch features, we tested whether the addition of pitch features to the onset, phonological feature, and peakRate model would improve model performance. Strikingly, we observed a clear separation between encoding of absolute pitch predominantly on the medial temporal plane (specifically medial HG and PT) and the encoding of relative pitch mostly in middle to anterior STG and in PP (Figure 4A, E). These observations were supported by a formal clustering analysis with pairwise comparisons of Silhouette index (Figure S4). Onset-encoding electrodes were confined to pSTG, with a clear separation from pitch encoding electrodes but not feature -encoding electrodes, which were also present in pSTG and extended anteriorly into midSTG (onset cluster pairwise permutation: p_relPitch_=0.001, p_absPitch_ = 0.001, p_feat_=0.14). Absolute pitch encoding populations were concentrated on the temporal plane and were anatomically separate from all other features (absolute pitch cluster pairwise permutation p_ons_=0.001 p_feat_=0.001, p_relPitch_=0.01), whereas feature and relative pitch encoding were both localized to middle STG with relative pitch encoding extending more anteriorly than feature encoding (pairwise permutation p_feat_=0.001, p_relPitch_=0.008). Notably, even though phonological features and relative pitch encoding were located within the same broad anatomical area, individual electrodes tended to encode one set of features at a time rather than the combination of the two. For example, 83% of electrodes that had significant feature encoding did not have significant relative pitch encoding (out of total 302 electrodes with significant feature encoding). For relative pitch electrodes, the same trend held, though less strongly -- 51% of relative pitch encoding electrodes did not significantly encode features (out of total 105 electrodes with significant relative pitch encoding).

### Uncovering temporal dynamics of functional areas via unsupervised clustering

The encoding models described above are a method of uncovering representations of specific features chosen from an *a priori* hypothesis of cortical function. However, we were also interested in understanding the specific temporal response properties within each region. In a previous study, we showed that the STG could be parcellated into two distinct regions -- a posterior region that was selective for acoustic onsets and responded in a strongly adaptive manner throughout a sentence, and a mid-to-anterior region that responded in a sustained manner, without strong adaptation over time (Hamilton et al. 2018). Here, we asked whether these same temporal response types might also define the main sources of variance in the temporal plane auditory cortex, using the same data-driven approach to cluster activity in primary and non-primary auditory cortex according to dominant temporal response profiles. This approach replicated our previous findings of spatially localized onset selectivity in pSTG, and provided important insight into the differences between temporal plane and lateral STG areas, including changes in response latency.

To find patterns of activity that were replicable across all participants, we applied convex non-negative matrix factorization (cNMF, see methods) to the average responses to speech stimuli from all responsive electrodes on the temporal plane and lateral STG. This method decomposes stimulus-evoked neural signals into component time series that best describe the dominant temporal response profiles.

Each electrode is assigned a cluster weight that describes how to reconstruct its activity from a linear combination of these component time series. By averaging across all sentences, we were able to capture response latency differences across cortex that may not be captured across disparately tuned sites. From this analysis, we found that responses to speech over the entire auditory cortex can be decomposed into 4 classes that explain 89% of the variance in average responses to sentences (Figure 5A, inset). These clusters differed in their response profiles across sentence time courses, in particular in their response peak latencies, as measured by the time of the peak of the average beta weight for our STRF models (Figure 6B, Kruskal-Wallis ANOVA p = 1e-14, df=3.00, chi-sq=68.19) Specifically, these temporal response profiles included (1) a fast (short-latency), mixed onset and sustained response mostly on the temporal plane (Figure 5A-C, green traces), and some STG; (2) fast (short-latency), onset only responses confined to pSTG only (Figure 5A-C, purple traces); (3) a slower, mixed onset+sustained response seen mostly in STG (Figure 5A-C, pink traces); and (4) a much slower sustained responses throughout sentences, confined to anterior temporal plane (PP) and STG (Figure 5A-C, orange traces). Responses of single electrodes from each of these clusters are shown in Figure 5D. Perhaps most strikingly, responses in the onset only cluster (purple) were confined to pSTG and not observed on the temporal plane. On the other hand, a separate cluster on the temporal plane and lateral STG (green) showed some increased response at onset, but a continued response throughout the sentence rather than a rapid return to baseline as in the pSTG onset region.

**Figure 5.**
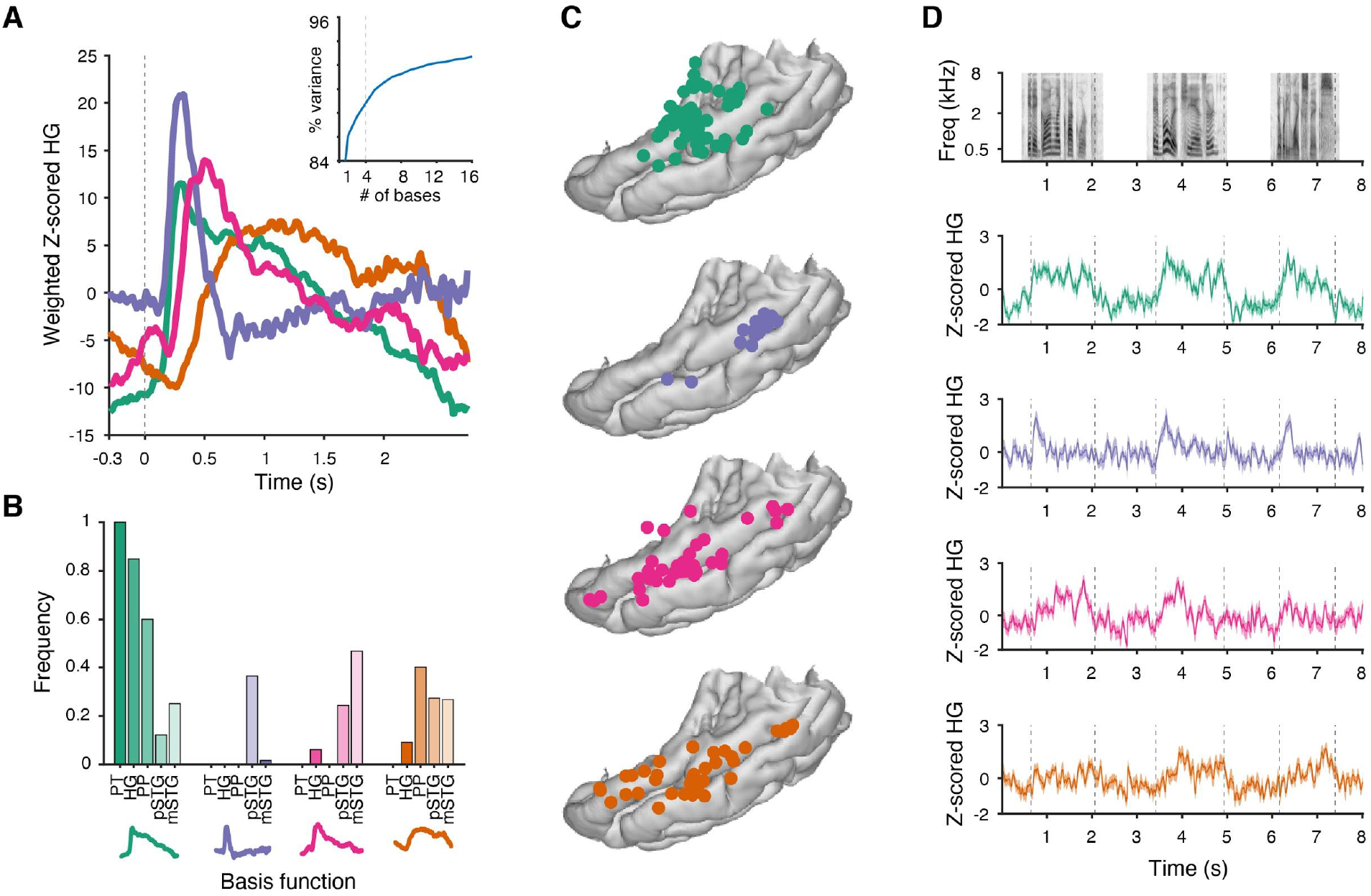
Unsupervised analysis of core and parabelt auditory cortex reveals posterior onset area is unique to pSTG and is active at latencies similar to auditory core (HG). (A) Canonical temporal response profiles to speech as measured through unsupervised cNMF. These Auditory cortical responses to speech could be decomposed into 4 classes that explain 89% of the variance in average responses to sentences (inset). These temporal response profiles included (1) a fast (short-latency), mixed onset+sustained response (green), (2) a fast (short-latency), onset only response (purple), (3) a slower, mixed onset+sustained response, and (4) a very slow sustained response. (B) Proportion of sites with each temporal response type within each area, e.g. sum across all bars for each area sum to 1 or less. (C) Anatomical distribution of sites in each of the four clusters. (D) Example single electrode traces from clusters 1 – 4.

**Figure 6.**
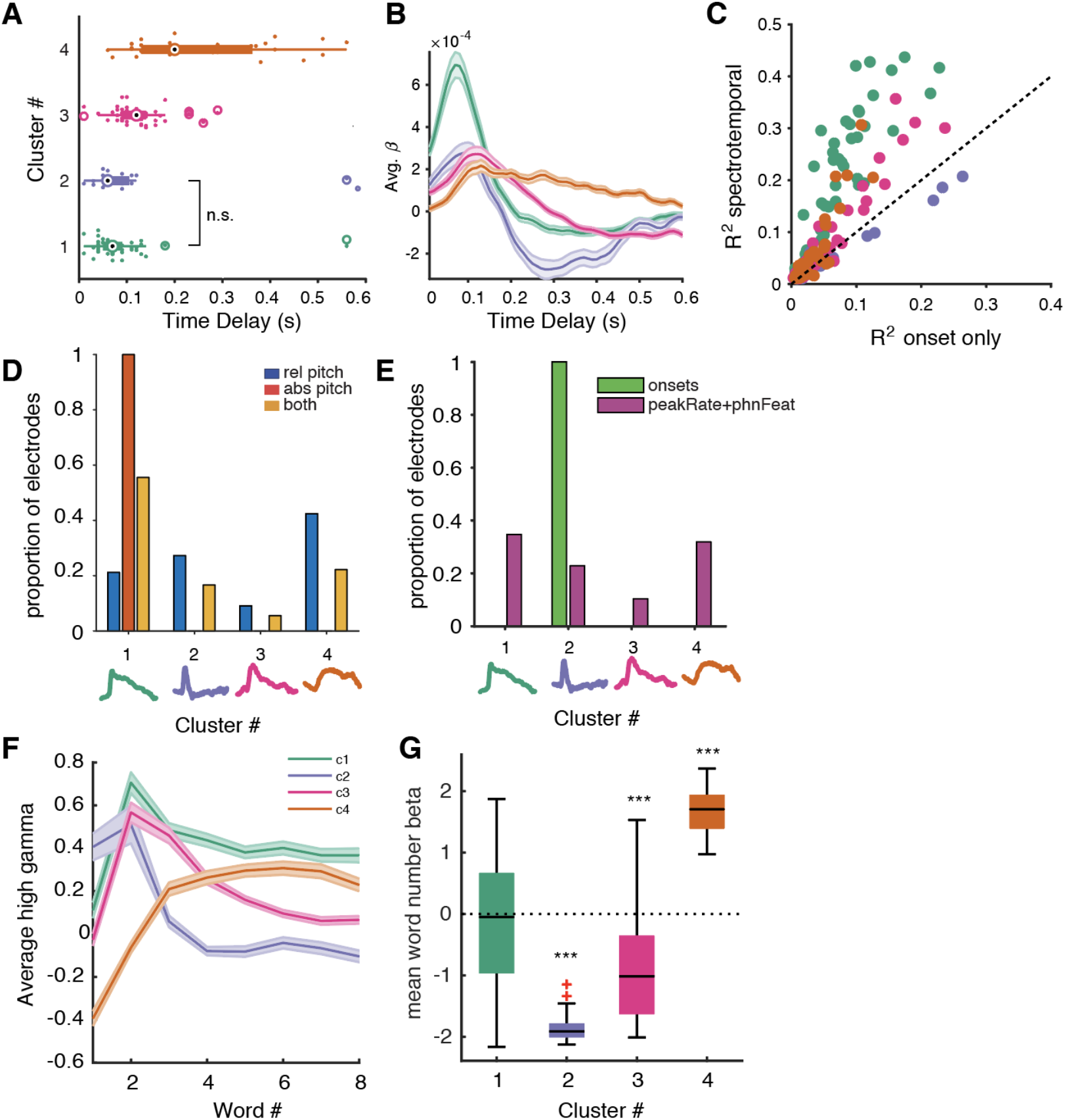
(A) The latency of the pSTG response is not significantly different from the latency of responses in temporal plane cluster 1. However, these two clusters are significantly faster to respond than cluster 3 and 4 (on STG and PP/STG respectively). Kruskal-Wallis ANOVA p = 1e-14, df=3.00, chi-sq=68.19 for effect of cluster. Post-hoc tests (Tukey-HSD corrected, comparing average rank): cluster 1 vs. 2 p=0.99, cluster 1 vs. 3 p=0.0001, 1 vs 4 p < 4e-9, cluster 2 vs 3 p=0.03, 2 vs 4 p = 7e-7, 3 vs 4 p = 0.0018. (B) Time course of average ß weights for the spectrotemporal model for each cluster. The peak of this function was used to calculate the latencies shown in (A). (C) Performance of a spectrotemporal model vs. an onset feature only model. A subset of electrodes in Cluster 2 clearly delineates a population that is best modeled by sentence onset. Electrodes in clusters 1, 3, and 4 are better modeled by the full spectrotemporal model rather than an onset-only model. (D) The proportion of electrodes in each cluster for which pitch predictors explained significant variance (permutation test). Absolute pitch dominates Cluster 1, whereas relative pitch is the main pitch feature in clusters 2 - 4. (E) Proportion of electrodes in each cluster for which onset or peakRate+phonological features explained significant variance (permutation test). 100% of electrodes in Cluster 2 had variance explained by the onset feature, whereas peakRate+phonological features explained variance in all clusters, with highest proportions in Clusters 1 (green) and 4 (orange), as was seen for lateral STG sites (Fig. 4F). (F) Average high gamma response increases with number of words in the sentence for Cluster 4 (orange), but not other clusters. Onset cluster 2 (purple) shows a large response during earlier words in the sentence. (G) Average beta weights for predicting cluster activity as a function of the number of words in the sentence. Cluster 4 electrodes show a strong positive relationship, whereby the number of total words is strongly positively predictive of a response, whereas Clusters 2 and 3 show a negative relationship. *** indicates p<0.001.

Pairwise comparisons of response latencies between single clusters showed that the response latency of the pSTG cluster 2 was not significantly different from the response latency in temporal plane cluster 1 (green, p=0.99, Figure 6A-B). However, these two clusters were significantly faster to respond than cluster 3 and 4 (pink and orange; on STG and PP/STG respectively, all p’s <0.05).

Our unsupervised clustering analysis clearly isolates the subset of pSTG electrodes that can be explained by a single sentence onset feature (Figure 6C – compare to Figure 4C, left panel, Figure 6E). In addition, it independently corroborates the finding of absolute pitch encoding electrodes appearing mostly on the temporal plane (Figure 6D). Absolute pitch electrodes were confined to cluster 1, whereas relative pitch electrodes were most frequently observed in cluster 4. Onset electrodes were confined to cluster 2, whereas features were distributed across clusters. The electrodes for which the onset predictor explained unique variance were confined to cluster 2, whereas peakRate and phonological feature representations were distributed across all clusters. This separation of features occurred despite the fact that clustering was performed on average responses, suggesting that temporal dynamics of these responses may also relate to their functional tuning.

The unsupervised clustering analysis isolated three distinct clusters on lateral STG, of which cluster 4 contained a large population of anterior lateral STG and PP electrodes and was characterized by slow delayed responses to sentence onsets. We hypothesized that this cluster might contain neural populations involved in higher order linguistic processing over the course of a sentence. Recent work (Fedorenko et al., 2016) identified such populations as having increased neural responses with additional words in a sentence. Indeed, we found that average neural response magnitude in cluster 4, but not in clusters 1-3, increased with word number in sentence (Figure 6F). Linear regression weights for the effect of word number on response magnitude within single electrodes differed significantly between clusters (F(3,147) = 90.8, p<0.001), with positive weights in cluster 4 (t(36) = 27.8. p<0.001) only. Cluster 1 showed no effect of word number on response magnitude (b = −0.1, p=0.5), while clusters 2 and 3 showed the opposite effect (cluster 2: b = −1.82, t(14) = −24.5, p < 0.001, cluster 3: b = −0.9, t(46) = −27.2, p <0.001, Figure 6G). As word number in the sentence is positively correlated with time from sentence onset, we further tested whether adding time from sentence onset to the model would abolish the effect of word number in cluster 4 using linear mixed-effects modeling. We found that adding word onset time to the model did not abolish the effect of word number (b = 0.03, SE = 0.01, t = 2.9, p = 0.004). This suggests that this effect is not due to delayed responses after sentence onset, but might reflect additional information processing as sentences unfold. Taken together, this analysis thus supports the notion of higher-order linguistic processing in anterior STG, in contrast to acoustic-phonetic speech feature representations in posterior and mid-STG.

Overall, our unsupervised clustering results provide additional support for a super-class of temporal response profiles (onset and sustained, or fast and slow) that may have additional speech feature selectivity (phonological, temporal or pitch) within this set. Because the analysis was performed on average responses to sentences rather than each sentence individually, we were able to capture the temporal dynamics of responses that were separate from tuning to individual speech features. Although some of these temporal clusters also coincide with specific feature representations (e.g. cluster 1 has strong absolute pitch representations), this is not a requirement of our analysis. Taken together, these results provide a detailed view of the temporal dynamics of speech representations in the human superior temporal gyrus and temporal plane.

## Discussion

The human auditory system decomposes the speech signal into components that are relevant for perception. In this study, we provide a full characterization of speech responses across the auditory cortex-- including the Heschl’s gyrus, surrounding areas of planum temporale and planum polare, and the lateral STG. Microsurgical access to the Sylvian fissure provided dense simultaneous recordings of the highly heterogeneous responses to speech from all regions of the human auditory cortex, in contrast to previous intracranial approaches that relied on piecemeal sampling from one of these regions at a time. This allowed us to assess functional and anatomical parcellations for speech representation. In particular, we assessed the spatial localization of onset, phonological feature, peakRate, absolute pitch, and relative pitch representations across auditory areas of the temporal plane and lateral STG. While some of these representations were anatomically localized (e.g. onset and absolute pitch), others were distributed across multiple anatomical regions. The extraction of acoustic and phonological features is represented by highly heterogenous pattern of local encoding at specific sites throughout auditory cortex.

In the posteromedial aspect of the temporal plane, including Heschl’s gyrus and planum temporale, we find strong responses to pure tones, simple single-peak frequency receptive fields, absolute pitch tracking in speech, and similar spectro-temporal receptive fields for tones and speech. These patterns are in line with previous studies that employed depth recording electrodes in Heschl’s gyrus (HG) to show that core auditory cortical areas show tonotopic organization, fast latency responses to click trains, and can track pitch changes in pure tones (Brugge et al., 2009; Griffiths et al., 2010; Howard et al., 1996; Steinschneider et al., 2014). In contrast, responses in PP were more similar to higher order areas of mid-to anterior STG, with different receptive fields for tones and speech, little frequency selectivity, and encoding of relative rather than absolute pitch in speech.

In lateral STG, neural responses to speech were stronger than to tones, and receptive fields were complex, without narrow spectral tuning. Overall, we found three functionally distinct populations in lateral STG. In addition to replicating the existence of an onset zone in pSTG (Hamilton et al., 2018), we find functionally distinct, but anatomically interleaved, populations in mid-STG representing different linguistically-relevant features in speech. These included phonological features, acoustic onset edges that cue syllables (Oganian and Chang, 2019), and relative pitch, which is the main cue to intonational prosody (Tang et al., 2017). While relative pitch was also represented in PP, peakRate and phonetic features are represented predominantly in lateral STG, supporting the unique role of lateral STG for speech processing. These lateral representations allow for joint encoding of phoneme identity via combinations of phonological features, and stress via peakRate and relative pitch representations.

Converging evidence from unsupervised clustering revealed that neural populations that represent speech features are distinct from those representing speech onsets (Hamilton et al., 2018), as well as from more anterior populations that appear to integrate information across words at the sentential level (Fedorenko et al., 2016). Critically, this distinction does not follow sulcal and gyral anatomical landmarks, as the feature-encoding populations in mid-STG covered both sides of the anatomically defined boundary between posterior and anterior STG.

Processing for absolute versus relative pitch was distinctly regionalized. In our previous work on pitch intonation encoding in the STG, absolute pitch responses were rare compared to relative pitch and phonological representations (Tang et al., 2017). Here we find that absolute pitch selectivity dominated in the temporal plane, mostly in HG and PT. Such absolute pitch sensitivity has been observed in nonhuman primates at the anterolateral border of the auditory core (Bendor and Wang, 2005, 2010) and fits well with the narrow, low spectral tuning for pure tones and speech vocal pitch in these areas. In contrast, relative pitch representations dominated in mid-anterior lateral STG and PP. This is also consistent with previous studies of pitch perception in the human auditory cortex, which have shown that sounds with pitch activate more of lateral HG than sounds without pitch, and that sounds with pitch variation also activate regions of PP and anterior STG (Patterson et al., 2002). This selectivity may also be related to voice-selective areas on the anterior temporal plane in macaque (Perrodin et al., 2011; Petkov et al., 2008, 2009).

Notably, our data contained several functional response profiles spanning multiple anatomical regions. First, several sites in PT were strongly pure tone selective, consistent with “core”-like responses. This may be due to sampling of sites that are on the border to HG, or may be due to the previously described functional heterogeneity of PT, which may include multiple subfields and has been found to respond to tones and music in addition to speech (Galaburda and Sanides, 1980; Griffiths and Warren, 2002; Hickok and Poeppel, 2007). On the other hand, while some electrodes in HG had clear single-peaked classical receptive fields for pure tone stimuli, some electrodes showed more complex, multi-peaked responses. This is in line with suggestions in previous imaging research that the auditory core is only confined to part of the HG, and the boundaries of this functional area vary widely across participants who may also have differences in underlying structure, including duplications of the Heschl’s gyrus (Brewer and Barton 2016; Da Costa et al. 2011). Finally, our results suggest that lateral STG can be subdivided into three functional areas, including a posterior onset, middle speech feature, and an anterior higher-order region that integrates speech information over time. Taken together, our results thus lend support to a shift away from considering anatomical landmarks as delineators of functional areas, and rather, suggest that a joint functional-anatomical parcellation is more informative (Glasser et al. 2016).

To date, very few studies have investigated auditory representations in the planum polare (PP). Here, we find that response profiles in PP are most similar to response profiles in more anterior portions of mSTG. Both areas favored relative over absolute pitch. Moreover, electrodes in PP and aSTG jointly contributed to a slowly responding functional cluster, where response magnitude increased during a sentence, further supporting higher level processing and integration, perhaps at the level of sentence comprehension (Fedorenko et al., 2016). Of note, the timescales of responses in PP tended to be much slower than those in HG and PT, further supporting different functional roles for these regions. Overall, PP and anterior portions of mSTG are functional very similar in our data, with respect to temporal dynamics of response profiles and functional selectivity. This suggests that the mid-to anterior representations typically recorded with lateral ECoG grids on the STG also extend medially to the temporal plane, including PP.

Thus far, we have described how specific acoustic and linguistic features are represented across the temporal plane and lateral STG. In our functional clustering analysis, differences in the temporal response properties of these regions allowed us to further speculate on information flow in the auditory system that is involved in perceiving speech (Brodbeck et al., 2018; Hickok and Poeppel, 2007; Jasmin et al., 2019; Rauschecker and Tian, 2000; Saenz and Langers, 2014). Our findings agree with previously reported differences in spectral selectivity between the temporal plane and lateral temporal areas (Da Costa et al., 2011; Hall et al., 2002; Leaver and Rauschecker, 2016; Saenz and Langers, 2014). However, other aspects of our results challenge current models of information flow in the auditory cortex.

For example, we find comparably short response latencies in HG/PT and posterior STG. However, the representational content of these areas was very different. A distinct region of the pSTG responded to onsets exclusively, with a broad spectral but narrow temporal response, a pattern that was not seen in temporal plane regions. Here, and in our previous work, we have observed that this region is not purely speech-selective, but responds to nonspeech and synthetic sound onsets as well (Hamilton et al., 2018). In contrast, HG/PT spectro-temporal receptive fields were more selective spectrally. The similar response timescales in these areas indicate that the onset zone in pSTG reflects early processing occurring in parallel to the computations performed by circuits on the temporal plane itself (Nourski et al., 2014). In line with this, a recent study of scans from the Human Connectome Project showed areas of higher myelination in posterior superior temporal gyrus that appear separate from the highest myelinated areas on Heschl’s gyrus and core auditory cortex (Glasser et al., 2014). Similar findings have also been observed by other groups investigating myelination of auditory cortex and nearby areas (Dick et al., 2012). Other groups have suggested that areas within the pSTG may be involved in processing multimodal information, including audiovisual information in speech (Ozker et al. 2017, Ozker et al. 2018). Overall, these results support the possibility of distributed parallel processing in medial temporal plane and posterior lateral STG for intermediate representations of speech including onset, phonological, and pitch information.

The independence of representations between temporal plane areas HG/PT and the pSTG onset area is also supported by clinical studies showing that resection of HG itself does not result in speech comprehension deficits (Russell and Golfinos, 2003; Sakurada et al., 2007; Silbergeld, 1997). This was reported even in tumor resections in the language dominant left hemisphere (Sakurada et al., 2007; Silbergeld, 1997). Additionally, consistent with our results on absolute pitch representations on the temporal plane, a case study reported that resection of HG (in this case on the right side) could result in transient amusia (Russell and Golfinos, 2003). In contrast, damage to left lateral STG results in severe speech comprehension deficits (Butler et al., 2014; Wernicke, 1874). Together, this suggests that the primary auditory cortex on HG is not the only source of inputs to the STG. We thus speculate that in addition to possible information flow from the caudal pSTG onset area to mid-anterior STG, the posterior and mid-anterior STG areas receive direct subcortical inputs that may bypass the auditory core.

While functional subfields of auditory cortex have been delineated in macaque models (Hackett et al., 2001; Kaas and Hackett, 2000), the existence of analogous subfields (for instance, a “belt” vs. “parabelt”) in humans is not agreed upon. “Core” auditory cortex is generally defined as the heavily myelinated, tonotopic region that receives its projections from the ventral MGB. “Belt” regions, on the other hand, surround the core and receive their projections from the core and dorsal MGB. The “parabelt” then receives its projections from the belt as well as the dorsal thalamus (Hackett et al., 2007; Rauschecker et al., 1997; Scott et al., 2017a, 2017b). While cytoarchitectonic distinctions between putative human core, belt, and parabelt areas have been demonstrated in some studies (Galaburda and Sanides, 1980; Sweet et al., 2005; Woods and Alain, 2009), the functional distinctions between these areas, particularly belt and parabelt, are less clear. Instead, many studies distinguish between “core” and “non-core” regions. For example, non-core regions are strongly modulated by attention (O’Sullivan et al., 2019) and invariant to background noise (Kell and McDermott, 2019), unlike core regions. Here, we provide evidence for distinct, parallel functional representations in the “core” and “non-core” regions, consistent with independent thalamocortical inputs.

Our results here illustrate both regionalized and distributed representations of the acoustic and linguistic features underlying the perception of human speech. Based on the differences in representation and the temporal latencies of these areas, we also suggest a refinement of dominant theories that posit a simple hierarchical flow of auditory information for speech processing from medial to lateral STG, or along the posterior to anterior axis of lateral STG. We find that speech representations are distributed across the human auditory cortex, with evidence of parallel processing on the temporal plane and in posterior STG for broadband onset and narrowband spectrotemporal information. We also argue that relative pitch, phonological features, and peakRate are intermediate levels of representation, computed in parallel. Overall, our findings speak to a distributed mosaic of specialized processing units, each representing different acoustic and phonetic cues in the speech signal, the combination of which creates the rich experience of natural speech comprehension.

## Methods

The University of California, San Francisco Institutional Review Board approved all procedures, and all patients provided written informed consent to participate.

We acquired data in an acute intraoperative setting from 9 patients (8 M/1 F, mean ± stdev. age: 32 ± 12 years) undergoing left hemisphere insular or opercular tumor resection (N=5) or phase II monitoring for intractable epilepsy (N=4). In patients with tumors, the location of their tumors near eloquent cortex necessitated dissection of the Sylvian fissure, which thereby provided access to the temporal plane. In epilepsy patients, temporal plane access was clinically indicated for seizure monitoring (seizure onset zone thought to originate from within Sylvian fissure). In 9 patients, we acquired simultaneous neural recordings directly from the temporal plane and the lateral surface of the brain (including the superior temporal gyrus and middle temporal gyrus). In 7 patients, 32 channel (8 x 4) or 64 channel (8 x 8) grids with 4mm center-to-center spacing and 1.17 mm diameter exposed contact lateral grids (Integra or AdTech) were placed on the temporal plane. In 8 patients, recordings from the lateral surface were acquired using grids with identical specifications (4-mm spacing and 1.17 mm diameter) but either 256 channels, 64 channels, or 32 channels total. In one patient, bilateral stereo EEG depth electrodes provided coverage of sites in Heschl’s gyrus. Patient demographics and the specifics of grid versus depth recording coverage are detailed in Table 1.

**Table 1.** Participant information, including location of electrodes, demographic information, handedness and language dominance, stimulus availability, and resection location.

| Participant | Hemisphere /<br>number elec | Age | Sex | Handedness | Language<br>Dominance | TIMIT | Pure<br>tone | Resection |
| --- | --- | --- | --- | --- | --- | --- | --- | --- |
| S01 | L, 98 | 52 | M | R | L | Y | N | Left insular<br>tumor |
| S02 | L, 54 | 29 | M | R | L | Y | N | Left frontal<br>operculum<br>tumor |
| S03 | L, 97 | 26 | M | R | L | Y | Y | Left frontal<br>operculum<br>tumor |
| S04 | L, 100 | 43 | M | R | L | Y | Y | Left insular<br>tumor |
| S05 | L, 50 | 35 | F | R | L | Y | N | Left fronto-<br>parietal<br>tumor |
| S06 | L, 90 | 21 | M | R | L | Y | N | Anterior<br>temporal<br>superior<br>temporal<br>gyrus |
| S07 | L, 79 | 19 | M | R | L | Y | Y | No<br>resection,<br>RNS<br>placed for<br>seizures in<br>left STG<br>and MTG |
| S08 | Bilateral<br>SEEG, 14 left | 43 | M | R | L | Y | Y | Left<br>temporal |
| S09 | L grid +<br>depths, 54 | 22 | M | R | L | Y | Y | Left<br>anterior<br>temporal<br>lobe |

**Table 2.** Post-hoc comparisons of response magnitude for pure tones and speech stimuli, grouped by auditory area. PT = planum temporale, HG = Heschl’s gyrus, PP = planum polare, pSTG = posterior superior temporal gyrus, mSTG = middle superior temporal gyrus. Multiple comparisons were calculated with MATLAB multcompare function, correcting for multiple comparisons using the Tukey HSD method.

| Post hoc tests (Tukey-Kramer) |  |  |  |  |
| --- | --- | --- | --- | --- |
|  | pure tones |  | speech |  |
| Contrast | Difference Estimate [range] | p-value | Difference Estimate [range] | p-value |
| PT vs. HG | 48.1, [-38.7, 135.0] | 0.555 | -47.26, [-134.11, 39.59] | 0.573 |
| PT vs. PP | 79.5, [3.4, 155.5] | 0.035* | -77.94, [-153.97, -1.90] | 0.04* |
| PT vs. pSTG | 109.1, [35.4, 182.9] | 0.0005*** | -127.82, [-201.55, -54.09] | <0.0001*** |
| PT vs. mSTG | 127.1, [52.3, 201.8] | <0.0001*** | -133.72, [-208.49, -58.94] | <0.0001*** |
| HG vs. PP | 31.4, [-28.9, 91.6] | 0.61 | -30.68, [-90.90, 29.54] | 0.634 |
| HG vs. pSTG | 61.0, [3.8, 118.3] | 0.03* | -80.56, [-137.84, -23.28] | 0.001** |
| HG vs. mSTG | 79.0, [20.3, 137.6] | 0.002** | -86.46, [-145.08, -27.84] | 0.0005*** |
| PP vs. pSTG | 29.7, [-9.3, 68.6] | 0.230 | -49.88, [-88.86, -10.91] | 0.0044** |
| PP vs. mSTG | 47.6, [6.7, 88.5] | 0.013* | -55.78, [-96.71, -14.86] | 0.0019** |
| pSTG vs. mSTG | 17.9 [-18.5, 54.38] | 0.665 | -5.90, [-42.35, 30.55] | 0.992 |

### Neural recordings

We acquired electrophysiological recordings at a sampling rate of 3051.8 Hz using a 256-channel PZ2 amplifier or 512-channel PZ5 amplifier connected to an RZ2 digital acquisition system (Tucker-Davis Technologies, Alachua, FL, USA). The local field potential was recorded from each electrode, notch-filtered at 60 Hz and harmonics (120 Hz and 180 Hz) to reduce line-noise related artifacts, and re-referenced to the common average across channels sharing the same connector to the preamplifier (Cheung et al., 2016). For STRF and clustering analyses, signals were bandpass filtered in the high gamma range (70-150 Hz) using the log-analytic amplitude of the Hilbert transform at 8 logarithmically-spaced center frequency bands within this range. We then took the first principal component across these 8 bands to extract stimulus-related neural activity (Edwards et al., 2009; Moses et al., 2016; Ray and Maunsell, 2011). Signals were subsequently downsampled to 100 Hz, then z-scored relative to the mean and standard deviation of activity across a recording block.

### Cortical surface extraction and electrode visualization

For recordings that were performed intraoperatively (N=5 participants), no CT scan was available, so we localized electrodes on each individual’s brain using intraoperative photographs of grid placement. For recordings performed in a chronic in-patient setting (N=4 participants), we co-registered the preoperative T1 to a postoperative CT scan and localized electrodes using in house software (Hamilton et al., 2017). Pial surface reconstructions were created from preoperative T1 MRI scans using Freesurfer. For visualization of electrode coordinates in MNI space, we performed nonlinear surface registration using a spherical sulcal-based alignment in Freesurfer, aligning to the cvs_avg35_inMNI152 template (Fischl et al., 1999). This nonlinear alignment ensures that electrodes on a gyrus in the participant’s native space remain on the same gyrus in the atlas space, but does not maintain the geometry of the grid.

### Stimuli

#### Speech stimuli

Participants listened passively to 499 sentences taken from the TIMIT acoustic-phonetic corpus (Garofolo et al., 1993) (286 males/116 female talkers from different regions of the United States of America). Each sentence was repeated once with 0.4 s of silence in between each sentence. In addition, a subset of 10 sentences (5 male, 5 female talkers) was repeated 10 times. Presentation of these stimuli was controlled using custom MATLAB software on a Windows laptop, and played through free-field speakers (Logitech). Sentences were presented in pseudo-random order.

#### Tone stimuli

A subset of participants (N=5) also listened to pure tone stimuli, generated as 80 mel-spaced frequencies from 74.5 Hz to 8 kHz (to match the spectrogram representations of the sentence stimuli). Each sine wave pure tone was 50 ms in duration with a 5-ms cosine ramp at the beginning and end of the tone. Pure tones were played at 3 intensity levels at 10 dB spacing, with the lowest intensity calibrated to be minimally audible in the hospital room. Each pure tone frequency/intensity pair was repeated 3 times, with jittered inter-stimulus intervals to minimize predictability of the stimulus (range 0.28 s minimum ISI – 0.5 s maximum ISI).

### Pure tone receptive fields

To calculate the pure tone receptive field, we first took the high gamma signals time-locked to the onset of each tone, and constructed a post-stimulus time histogram (PSTH) from 0 (tone onset) to 500 ms, collapsing across repetitions. Classical receptive fields were constructed by calculating the average high gamma response to each frequency-intensity pair in a window defined by the peak of the PSTH. The magnitude of the pure tone response was calculated by collapsing across all frequency-intensity pairs to get the maximum Z-scored high gamma response for each electrode. As a proxy for “clean”/significant receptive field tuning, we created a binary mask (3 intensity bins x 80 frequencies) for each receptive field using the normalized tuning curve that was then rescaled from 0 to 3 (for the 3 intensity bins), and rounded to the nearest integer value. These amplitudes were then used to create a mask of 1s inside the receptive field (for all frequencies where the tuning curve amplitude met a given intensity value bin). This procedure effectively creates a matrix of NaNs for frequencies in the “background” (outside of the classical receptive field) and a (typically V-shaped) set of 1s for frequencies inside the receptive field. To determine whether sites showed significant tuning in this way, we calculated the amplitude of responses to tones of each frequency and intensity outside the receptive field and compared them to responses to tones at each specific frequency and intensity inside the receptive field. We then used a Wilcoxon rank sum test to calculate the difference in mean response for tones inside versus outside the receptive field. Receptive fields with “significant” tuning are shown with red axes in Fig S2 and S3, where it is clear that the response inside the tuning curve has a higher amplitude (stronger purple values) compared to tones outside the receptive field. For non-significant sites, overall amplitudes to tones may be greater than silence, but show no strong tuning to a given frequency range.

To calculate whether responses to tones were significantly greater than silence, we computed a Wilcoxon rank sum test to assess the difference in activity for 50 ms of pre-trial silence and tone-evoked activity from 0-250 ms following tone onset. P-values were corrected for multiple comparisons using the Bonferroni method, where the number of comparisons was equal to the number of electrodes being evaluated.

### Linear receptive field analysis

To uncover tuning for spectro-temporal and acoustic-phonetic features in individual electrode sites, we fit linear receptive field models (Aertsen and Johannesma, 1981; Theunissen et al., 2001) of the form:

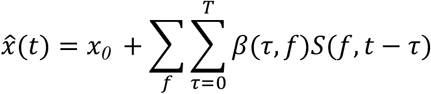

Where x is the neural activity recorded at a single electrode, *β*(*f*, *τ*) contains the regression weights for each feature *f* at time lag *τ*, and *S* is the stimulus representation for feature *f* at time *t* − *τ*. We also fit an intercept x_0_ for each electrode to allow for differences in baseline activity. We included delays of up to 600 ms to model longer latency responses that may be observed in mid-to anterior-STG sites (Hamilton et al., 2018). *β* weights were fit using ridge regression. The ridge parameter was estimated using a bootstrap procedure in which the training set was randomly divided into 80% prediction and 20% ridge testing sets. The ridge parameter was chosen from a range of 30 log-spaced values from 10^−2^ to 10^7^, as well as a ridge parameter of 0 (no regularization). The final value was chosen as the parameter that gave the best average performance across electrodes as assessed by correlation between the predicted and ridge test set performance. Once an optimal regularization parameter was chosen, the model was then trained on the complete training set. The final performance of the model was computed on a final held out set not included in the ridge parameter selection. Performance was measured as the correlation between the predicted response on the model and the actual high gamma measured for sentences in the test set.

We estimated models using a mixture of stimulus representations, including a mel-band spectrogram, sentence onset features (Hamilton et al., 2018), phonological features (Mesgarani et al., 2014), absolute and relative pitch features (Tang et al., 2017), and peakRate, calculated as the maximum change in the derivative of the acoustic envelope (Oganian and Chang, 2019). These features are described below and were used in combination to estimate the unique amount of variance explained by each feature (for example, a full feature model included sentence onset, peakRate, phonological features, absolute and relative pitch features).

#### Mel-band spectrogram

The spectrograms of each sentence were calculated using a mel-band auditory filterbank of 80 filters with center frequencies from approximately 75 Hz to 8 kHz. This frequency decomposition is thought to reflect the filtering performed by the human auditory system (Slaney, 1998), and has been used extensively in our previous work to describe spectro-temporal tuning within the STG (Hamilton et al., 2018; Mesgarani et al., 2014).

#### Sentence onset feature

The sentence onset feature consisted of a binary vector of values, with a 1 at the onset of the first sample of the first phoneme of each sentence, and 0s elsewhere.

#### Peak rate features

Peak rate was calculated as local peaks in the derivative of the amplitude envelope of speech (Oganian and Chang, 2019). First, we extracted the amplitude envelope of speech using the specific loudness method by Schotola (Schotola, 1984). This method first decomposes the speech signal into critical bands based on the Bark scale. Signals were square-rectified within each filter bank, bandpass filtered between 1 and 10 Hz, downsampled to 100 Hz, and then averaged across frequency bands to get the envelope. We then calculated the derivative of this envelope, and extracted local peaks in this derivative to create a sparse time series of “peakRate” features.

#### Phonological features

Phonological features consisted of binary phonological features used in our previous work (Hamilton et al., 2018; Mesgarani et al., 2014). These features describe single phonemes as a combination of voicing, place and manner of articulation features. They are a reduced representation of the speech sound signal that better captures responses to speech in non-primary auditory cortex (Mesgarani et al., 2014). As with sentence onset features, these matrices include a 1 at the onset of each phonological feature, and a 0 elsewhere. Features included sonorant, obstruent, voiced, back, front, low, high, dorsal, coronal, labial, syllabic, plosive, fricative, and nasal.

#### Absolute pitch features

Absolute pitch was calculated using procedures identical to (Tang et al., 2017). In brief, the fundamental frequency (F0) was calculated using an automated autocorrelation method in Praat, and manually corrected for doubling or halving errors. Absolute pitch was calculated as the natural logarithm of pitch values in Hz. We then created a binary feature matrix by discretizing these pitch values into 10 bins, equally spaced from the 2.5 to the 97.5 percentile values.

#### Relative pitch features

Relative pitch was also calculated using procedures identical to (Tang et al., 2017). The fundamental frequency (F0) was extracted as described above for absolute pitch. Relative pitch was calculated by z-scoring the log-F0 absolute pitch values within each sentence (within speaker), such that values were high or low relative to the average pitch of the speaker of that sentence. These normalized values were then discretized into 10 bins, as above. To calculate relative pitch change, we took the derivative of the z-scored log-F0 relative pitch values, and then discretized this pitch derivative curve.

#### Variance portioning to determine unique variance explained

To calculate the unique variance explained by a given set of features, we calculated R^2^ values for a reduced model that did not include these features of interest but did include other confounding features. We then compared these to R^2^ values to an extended model including the features of interest and the features from the reduced model. For example, to calculate the unique variance explained by phonetic features and peakRate together, we compared a model including onsets, phonetic features and peakRate to a model including onsets only. We used the following combinations of models to determine unique R2 values:

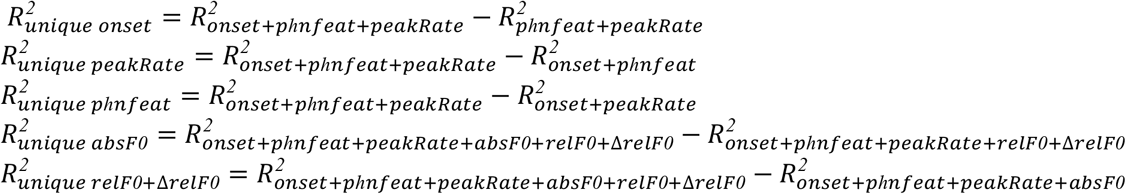

To determine whether the addition of a feature or set of features resulted in a statistically significant increase in R^2^, we used permutation testing. In these tests, the added feature labels were shuffled in 2.4-second chunks over time to create a shuffled distribution of feature values. 2.4 seconds was chosen as 4 times the length of the delay period of 0.6 seconds. We then calculated the change in R^2^ that was observed by adding these (dummy) variables to the full model. Each feature shuffle was performed 1000 times, allowing us to determine significant differences up to a threshold of p<0.001.

#### Calculating frequency tuning curves for speech

To calculate frequency tuning curves for speech and compare them to pure tone responses, we took the speech-derived STRF (using the mel-band spectrogram), summed across all time delays, and smoothed the resulting frequency tuning curve using a 3rd-order Savitzky-Golay filter with a 31-point window. Example frequency tuning curves for speech and pure tones are shown in Fig. 2E. The number of peaks in each tuning curve was calculated using the findpeaks function of the MATLAB signal processing toolbox on each tuning curve normalized to its maximum, with a minimum peak prominence of 0.5 and and a minimum peak height of 0.5.

To calculate the correlation between pure tone and frequency tuning (Fig. 2H), we used the frequency tuning curves defined above, and calculated the Pearson correlation between these tuning curves for each electrode. In Fig. 2H, only correlations where at least one stimulus type (pure tones or speech) elicited a significant response were included.

### Unsupervised clustering of local field potential time series

We performed unsupervised clustering of electrodes in the superior temporal gyrus, middle temporal gyrus, and temporal plane (planum temporale, Heschl’s gyrus, and planum polare) across all participants using convex non-negative matrix factorization (cNMF), similar to (Hamilton et al., 2018). In brief, cNMF uses an iterative decomposition to estimate the neural high gamma time series X [n time points x p electrodes], according to the following equation:

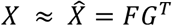

Where

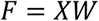

The G matrix [p electrodes x k clusters] represents the spatial weight of an electrode in a given cluster, and W [p electrodes x k clusters] represents the weights applied to the electrode time series. In order to look for canonical response types across the whole auditory cortex, we performed NMF on the average responses across the sentence stimuli that were heard by all participants. To choose the number of basis functions, we calculated the percent variance explained by each number of basis functions from k=1 to k=16, and chose the number of clusters at the elbow of the percent variance curve, resulting in a final number of k=4 clusters. These clusters represented approximately 89% of the variance in auditory cortical responses.

Although the NMF weights *G* are continuous and allow electrodes to belong to more than one cluster, for some analyses we assigned electrodes to their “best” NMF cluster. This was performed by sorting the NMF spatial weights within each electrode, and finding electrodes where the maximum cluster weight on the 4 clusters was greater than two times the next highest weight, and where the highest spatial weight was at least 0.1 (to avoid assigning electrodes to a cluster where no particular temporal response profile was a good match).

## Acknowledgments

The authors would like to thank Matthew Leonard, Patrick Hullett, and Brian Malone for helpful comments on the manuscript. In addition, the authors would like to thank Neal Fox, Matthew Leonard, Matthias Sjerps, Kunal Raygor, and Leah Muller for assistance with intraoperative recordings. This work was supported by grants from the NIH (F32 DC014192-01 Ruth L. Kirschstein postdoctoral fellowship, to LSH, and DP2-OD00862 and R01-DC012379 to EFC). E.F.C is a New York Stem Cell Foundation-Robertson Investigator. This research was also supported by The New York Stem Cell Foundation, The McKnight Foundation, The Shurl and Kay Curci Foundation, and The William K. Bowes Foundation. We gratefully acknowledge the support of NVIDIA Corporation with the donation of the Tesla K40 GPU used for this research. LSH and EFC conceived of and designed the experiment. LSH, YO, EFC, and others collected the data. LSH and YO analyzed the data. EC performed surgery and grid implantation. LSH, YO, and EFC wrote the paper.

## Supplemental Figures

**Figure S1.**
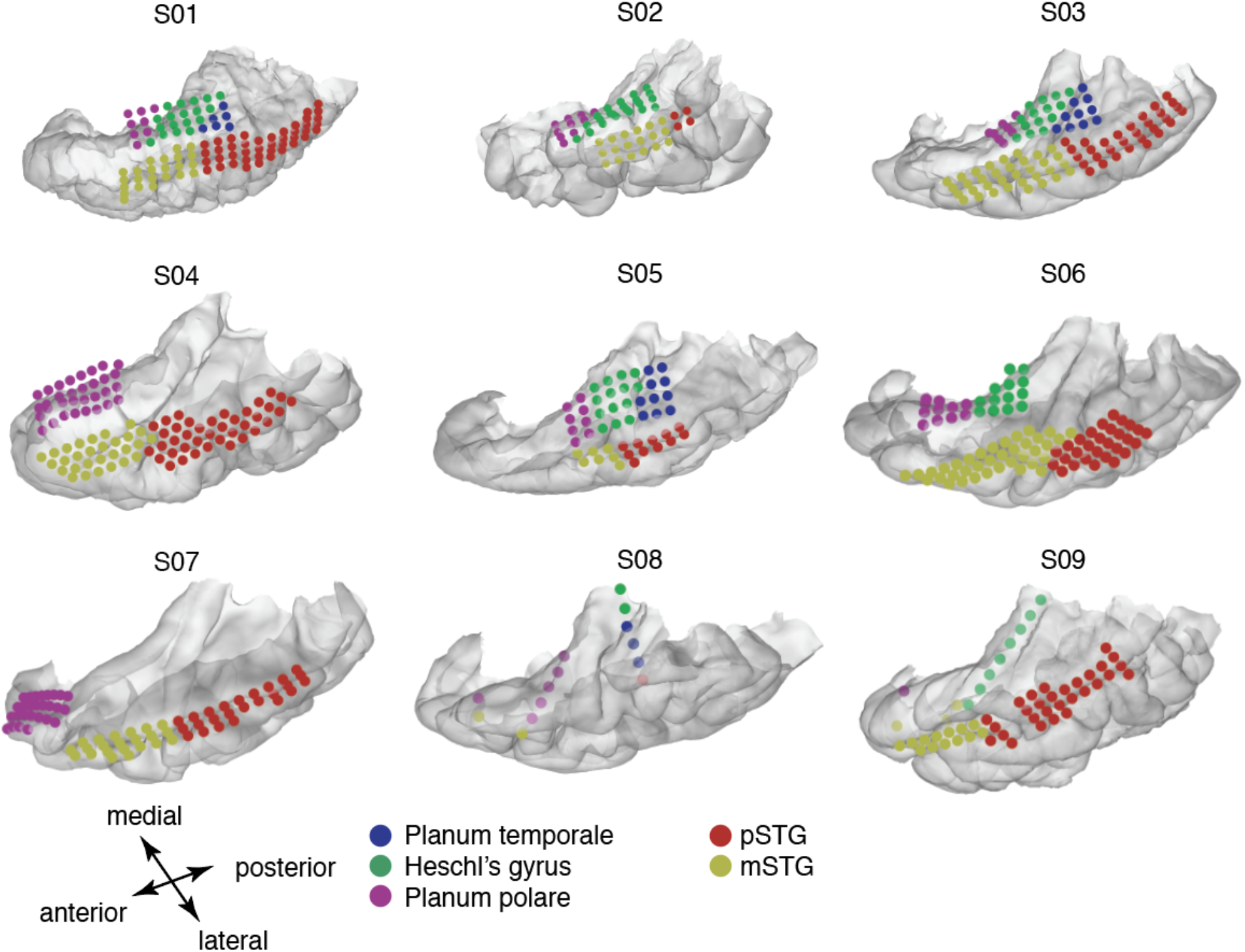
Electrodes from all participants across all auditory areas, shown on each participant’s left hemisphere temporal lobe reconstruction. Electrodes are colored according to anatomical region.

**Figure S2.**
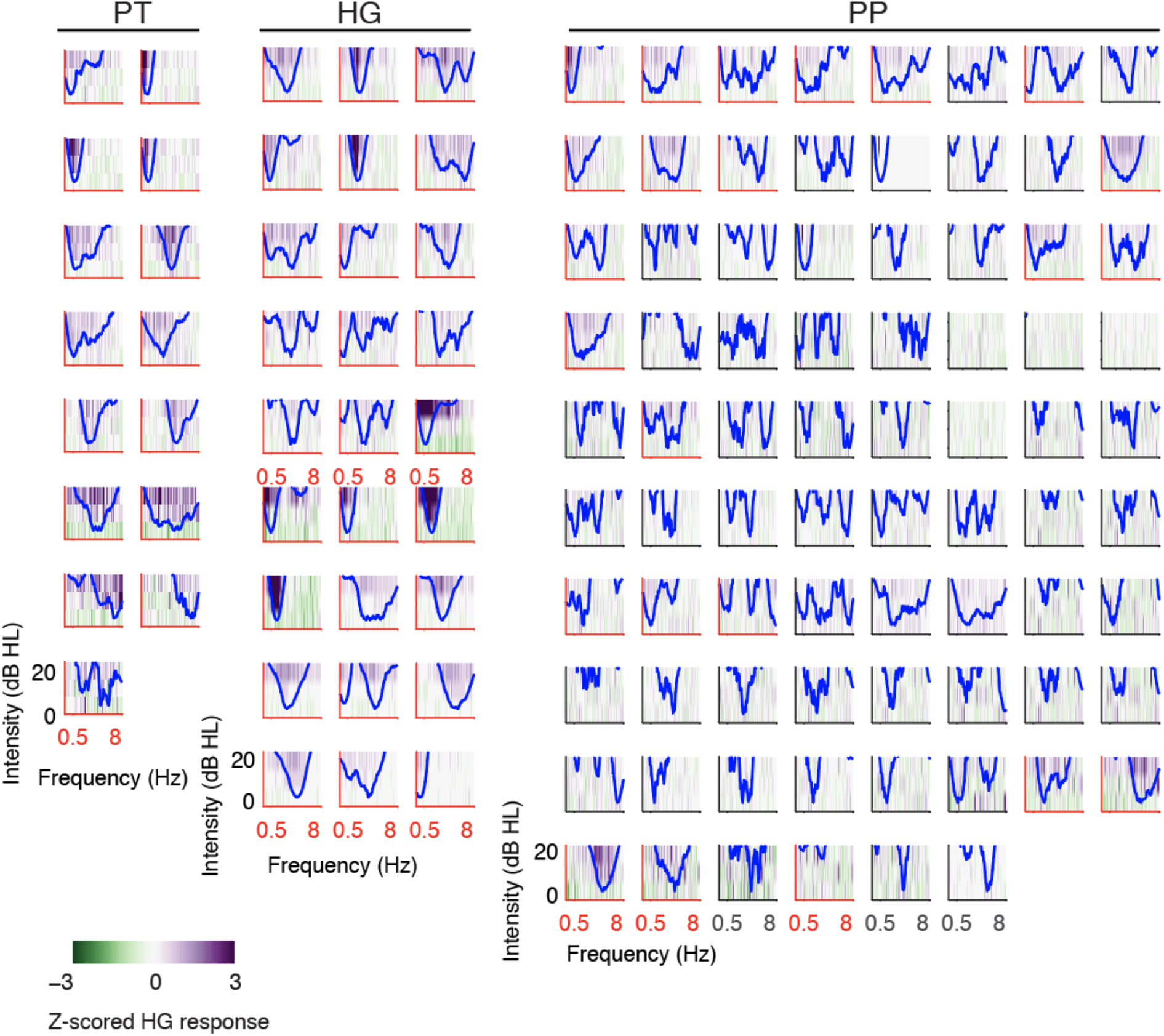
Pure tone receptive fields recorded from individual electrodes in PT, HG, and PP. Blue trace shows computed tuning curve based on smoothed receptive field data. Red axes indicate electrodes for which within receptive field responses were significantly larger (Bonferroni corrected p<0.05) than outside receptive field responses (i.e. inside or outside the blue tuning curve). This provides a proxy for “clean” receptive fields. Overall, most sites in PT and HG showed strong, classical V-shaped tuning for pure tones. PP sites tended to show multiple peaks or weak tuning.

**Figure S3.**
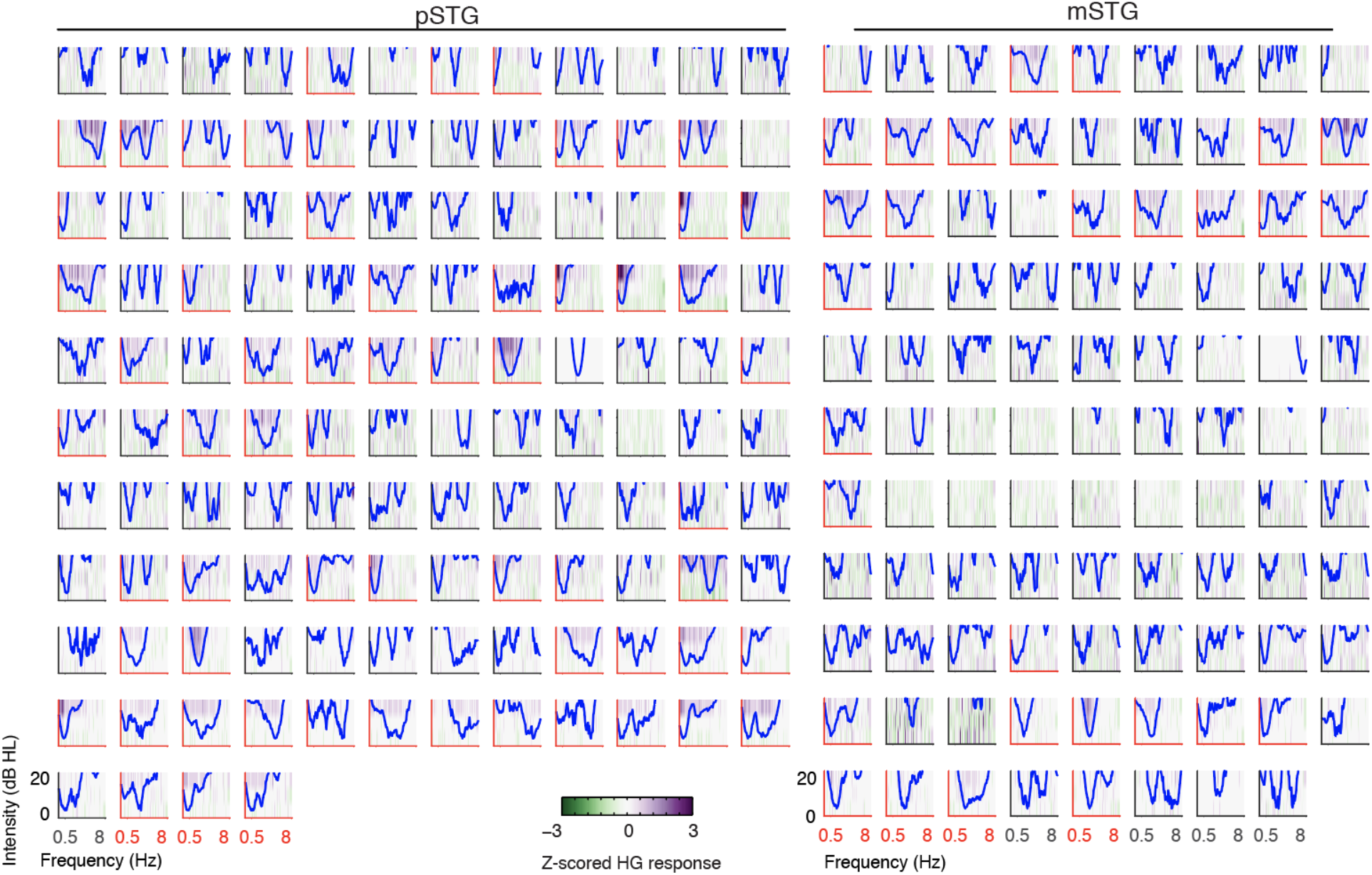
Pure tone receptive fields (RFs) recorded from individual electrodes in the posterior STG (pSTG) and mid-to-anterior STG (mSTG). Blue trace shows computed tuning curve based on smoothed receptive field data. Red axes indicate electrodes for which within receptive field responses were significantly larger (Bonferroni corrected p<0.05) than outside receptive field responses (i.e. inside or outside the blue tuning curve). This provides a proxy for “clean” receptive fields. Overall, sites in pSTG showed some pure tone tuning, but as shown in Figure 1, the magnitude of responses was overall lower compared to HG and PT. While some narrow-band, v-shaped tuning curves were observed in pSTG, many responses tended to be broader or multi-peaked. In mid-to-anterior STG, fewer sites with classical RFs were observed.

**Figure S4.**
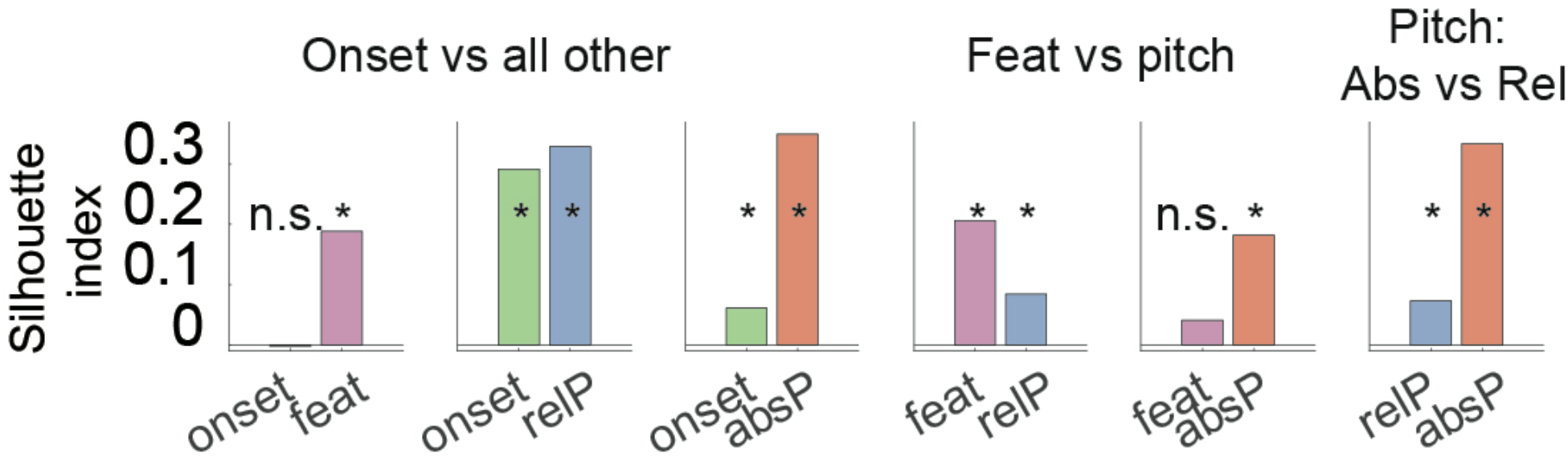
Anatomical clustering of electrodes with different feature encoding profiles. (A) Silhouette index for clustering of different encoding populations. Onsets cluster in pSTG, Phonetic features and relative pitch in mid-STG, absolute pitch on the temporal plane (^ p = .1, * p<.01, median permutation test).

## Notes

### Competing Interest Statement

The authors have declared no competing interest.

